# Sex differences in colonic inflammation are driven by epithelial-specific expression of estrogen receptor alpha

**DOI:** 10.1101/2024.09.28.615598

**Authors:** Guillermo A. Pereda, Adrian D. Kocinski, Alyssia V. Broncano, Sarah K. McNeer, Michelle L. Raymond, Nicholas P. Ziats, Keith A. Breau, Joseph Burclaff, Scott T. Magness, Wendy A. Goodman

**Affiliations:** Department of Pathology, Case Western Reserve University School of Medicine, Cleveland, Ohio, USA; Department of Cell Biology and Physiology, University of North Carolina at Chapel Hill, Chapel Hill, North Carolina, USA; Joint Department of Biomedical Engineering, University of North Carolina at Chapel Hill and North Carolina State University, Chapel Hill, North Carolina, USA; Center for Gastrointestinal Biology and Disease, University of North Carolina at Chapel Hill, Chapel Hill, North Carolina, USA; School of Medicine, University of North Carolina at Chapel Hill, Chapel Hill, North Carolina, USA

**Keywords:** Estrogen signaling, intestinal epithelium, inflammatory bowel disease, sex differences

## Abstract

**Background & Aims:** Inflammatory bowel disease (IBD) patients exhibit altered expression of nuclear estrogen receptors alpha and beta (ERα, ERβ) and G-protein coupled estrogen receptor 1 (GPER1). We previously showed that deletion of ERα protects against intestinal damage selectively in female mice; however, the mechanisms conferring sex-specific protection are poorly understood. The goal of this study was to compare ERα- and ERβ-specific mechanisms contributing to intestinal epithelial function in males and females.

**Methods:** Expression of ERα, ERβ, and GPER1 was evaluated in colonocytes from wild-type (WT) male and female mice. Intestinal epithelial cell (IEC)-specific ERα and ERβ knockout mice were developed and challenged with dextran sulfate sodium (DSS). Colonic organoids were used to identify estrogen-dependent and -independent effects on cellular growth, differentiation, and transcriptional regulation in WT, ERα-KO, and ERβ-KO IECs.

**Results:** Colonic IECs showed significant expression of ERα, ERβ, and GPER1 as well as Cyp19A1, which catalyzes production of 17β-estradiol (estrogen). Female mice lacking ERα specifically in colonic IECs showed protection from DSS-induced injury, whereas males showed increased pathology. Organoids derived from male ERα-KO mice showed enhanced proliferation and decreased expression of key functional genes even without exogenous estrogen; however, colonoids derived from female ERα-KO mice transcriptional analysis showed a protective gene signature. These findings reveal that deletion of ERα differentially contributes to enhanced barrier function and resistance to inflammation in females, but to dysfunctional hyper-proliferation in males.

**Conclusions:** ERα signaling within IECs drives opposing sex-dependent effects on the development, regenerative capacity, and inflammatory susceptibility of the intestinal epithelium.

## INTRODUCTION

Inflammatory bowel diseases (IBD), comprised of Crohn’s disease (CD) and ulcerative colitis (UC), are chronic inflammatory conditions of the gastrointestinal (GI) tract often resulting in comorbidities and severe disability (1). Although the clinical manifestations of CD and UC are distinct, both conditions result in a chronic, relapsing-remitting disease course that has no cure. The underlying pathophysiology of IBD is incompletely understood but thought to involve aberrant immune responses to intestinal flora in genetically susceptible individuals. Additional environmental factors, such as diet, smoking, and hormonal changes, also contribute to disease susceptibility. The prevalence of IBD has increased significantly over the past several decades, with more than 0.7% of Americans (2) and approximately 4.9 million individuals worldwide (3) currently diagnosed.

CD and UC exhibit distinct patterns of incidence, prevalence, and severity in males versus females (4). Large-scale meta-analyses of Western populations have identified an increased risk of CD among females beginning around the time of puberty, whereas adult males are at greater risk for UC (5). There are many contributing factors to sex differences in IBD, including genetics, the impact of sex hormones such as estrogens and androgens, and other environmental and social factors that differ among men and women. Changes in disease prevalence and severity around times of endocrine transition, such as puberty, pregnancy, and menopause, highlight the contributions of hormone signaling to intestinal inflammation. In particular, there is growing recognition that 17β-estradiol (estrogen, “E2”) exerts powerful effects on the immune system, modulating innate and adaptive immune responses (6, 7) and intestinal barrier function (8). However, the precise mechanisms by which E2 regulates tissue-level inflammation, including in the intestine, are incompletely understood.

E2 signals through three receptors, including estrogen receptors alpha and beta (ERα, ERβ) and a transmembrane receptor (GPER1). ERα and ERβ are nuclear receptors mediating gene transcription in target cells, and GPER1 is a G-protein coupled receptor mediating rapid signaling and second messenger activation in response to E2. All three receptors are broadly expressed outside of the reproductive system and mediate signaling in response to very low (pM) levels of E2 present in both males and females (9). Over 600 protein-coding genes express bona fide estrogen response elements (EREs) in their promoter regions, with potentially over 1000 additional genes predicted to be regulated by ER co-activation and/or co-repression (10). Mice lacking global expression of ERβ (ERβ-KO) exhibit changes to their intestinal morphology, including enhanced proliferation and impaired differentiation of colonic intestinal epithelial cells (IECs) (11). Similarly, the prostate epithelium of ERβ-KO mice is characterized by hyperproliferation and accumulation of incompletely-differentiated cells (11), suggesting that ERβ has an important role in epithelial proliferation and differentiation. Colonic expression of *Esr2* (encoding ERβ) is reduced in animal models and human IBD samples (8), and ERβ-specific agonist treatment was shown to improve disease in an *H. hepaticus* model of IBD (12). These observations, together with data showing increased epithelial monolayer resistance in trans-epithelial electrical resistance (TEER) assays (8), collectively suggest that expression and activation of ERβ helps to maintain colonic homeostasis.

Our previous work examined the relative protective contributions of ERα versus ERβ in dextran sulfate sodium (DSS)-induced intestinal injury in mice. Male and female ERβ-KO mice showed similar body weight loss and histological inflammation as age- and sex-matched wild-type (WT) controls (13), suggesting that loss of ERβ does not exacerbate inflammation *in vivo* in response to DSS. In contrast, ERα-KO mice showed sex-specific protection from DSS. Female ERα-KO mice maintained their body weight and showed minimal signs of colonic inflammation, whereas male ERα-KO mice that exhibited more severe weight loss and histological inflammation (13). Collectively, these data suggest that loss of ERα is protective in females.

The current study is focused on mechanisms underlying female-specific protection from intestinal injury in ERα-KO mice. We hypothesized that ERα may be a previously unrecognized but important regulator of epithelial function in the colon, contributing to sex-specific differences in ERα-KO mice. We undertook a systematic evaluation of estrogen receptor expression in primary colonocytes of male and female WT mice, finding that ERα is expressed at levels surpassing those of ERβ or GPER1. We developed novel mouse models lacking expression of ERα or ERβ specifically in intestinal epithelial cells and found that female-specific protection from DSS was recapitulated in ERα-KO conditional knockouts. Studies using *ex vivo* colonoid models revealed sex-specific differences in response to ERα deletion, including distinct transcriptional profiles in male versus female cells. Collectively, our findings identify ERα as a potent driver of sex-specific differences to colonic epithelial function, likely contributing to sex differences in response to intestinal injury and IBD (4, 5, 14, 15).

## RESULTS

### Colonic epithelial cells express significant levels of ERα and ERβ

We previously discovered that global deletion of estrogen receptor alpha (ERα) is protective against dextran sulfate sodium (DSS)-induced intestinal injury in female but not male mice (13). However, the interpretation of this finding is complicated by the broad expression profile of ERα outside of the female reproductive tract, including endothelial cells (16), adipose tissue (17), immune cells (18), and mammary gland epithelium (19). Importantly, several previous studies have shown strong expression of ERβ in colonic intestinal epithelial cells (IECs) of humans (20) and mice (11), where it is thought to promote barrier function and intestinal homeostasis (8), yet it is unclear if there are sex differences in IEC-specific expression of ERβ, or whether colonic IECs even express ERα. Therefore, we assessed the relative expression of ERα and ERβ at mRNA and protein levels in primary colonocytes isolated from healthy wild type (WT) mice (Figure 1).

**Figure 1:**
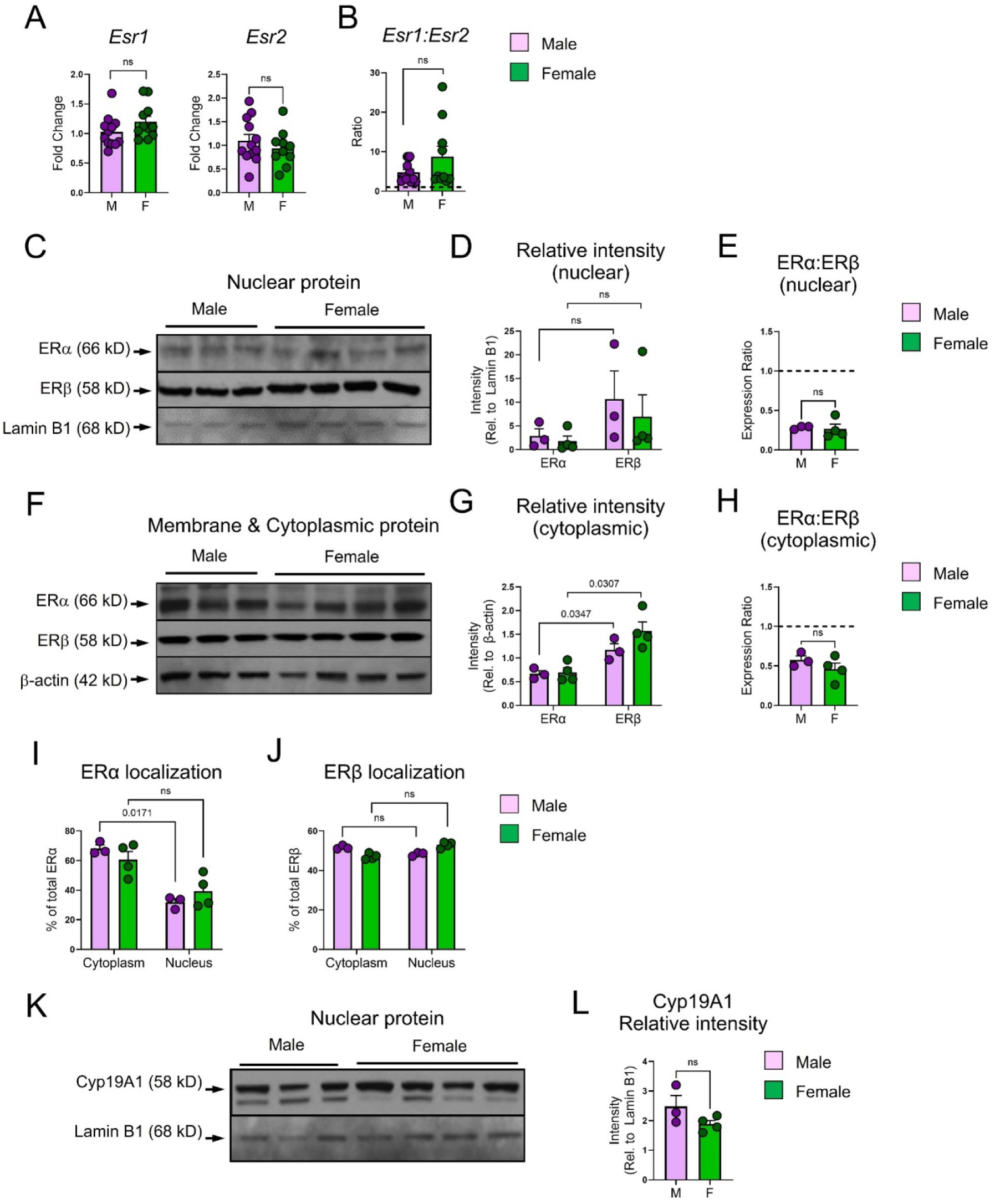
Colonic epithelial cells express significant levels of ERα and ERβ. *(A)* Relative gene expression of *Esr1* (ERα) and *Esr2* (ERβ) in primary colonic epithelial cells obtained from healthy adult wild-type (WT) mice was determined by qPCR. Expression of target genes was normalized to that of *Gapdh* and fold change was calculated relative to male samples. *(B)* Ratio of *Esr1:Esr2* mRNA expression was calculated for male and female samples. *(C-H)* Protein expression of ERα and ERβ were determined by western blot of nuclear *(C-E)* or cytoplasmic *(F-H)* protein lysates isolated from primary colonic epithelial cells of healthy adult WT mice. Expression of ERα and ERβ were normalized to levels of Lamin B1 (nuclear lysates, *(D)*) or β-actin (cytoplasmic lysates, *(G)*) using densitometry. Densitometry values were used to calculate ratios of ERα:ERβ expressed in nucleus *(E)* and cytoplasm *(H)*. Subcellular localization of ERα *(I)* and ERβ *(J)* was calculated using densitometry values. *(K)* Expression of Cyp19A1 (aromatase) was determined by western blot of nuclear protein lysates isolated from primary colonic epithelial cells of healthy adult WT mice and *(L)* quantified by densitometry. For all figures, statistical analysis was performed with 2-way analysis of variance (ANOVA) and Tukey post hoc test. Individual points represent individual animals.

We measured the transcription of *Esr1* (encoding ERα) and *Esr2* (encoding ERβ) using quantitative PCR (qPCR) and found comparable expression of *Esr1* and *Esr2* among male and female colonocytes (Figure 1A), with *Esr1* expressed at higher levels than *Esr2* in all samples (Ct values of 30-31 for *Esr1* versus 32-33 for *Esr2*). This revealed a 5 to 10-fold higher transcription of the *Esr1* gene compared to *Esr2*, with only modest sex differences observed (Figure 1B). However, in sharp contrast to transcript levels, we observed that protein levels of ERα were significantly lower than those of ERβ within both nuclear (Figures 1C-E) and cytoplasmic/membrane (Figure 1F-H) lysates. Analysis of the proportion of total ERα protein found in nucleus versus cytoplasm/membrane for each sample revealed that the majority of ERα is confined to the nucleus in both male and female IECs (Figure 1I). In contrast, ERβ is expressed equivalently in nucleus and cytoplasm (Figure 1J). These data demonstrate that expression of ERα and ERβ is predominantly under translational control.

We also assessed expression of Gper1, the membrane receptor for estrogen (21). Both *Gper1* mRNA (Supplemental Figure S1A-C) and protein (Supplemental Figure S1D-G) were found to be strongly expressed in murine IECs, with no significant differences in expression between male and female samples. Analysis of a publicly available single-cell RNA-Seq dataset (22) further revealed that in human samples, *GPER1* is significantly higher in female colonocytes than males (Supplemental Figure S1H) and is more strongly expressed in the distal compared to proximal colon (Supplemental Figure S1I).

Lastly, we analyzed colonic IECs for the capacity to produce estrogen from cholesterol precursors by assessing the expression of Cyp19A1, the aromatase that catalyzes the final step of estrogen biosynthesis. We found high expression of Cyp19A1 in nuclear IEC lysates from male and female WT mice (Figure 1K-L), indicating that IECs are capable of synthesizing estrogen.

Collectively, these results demonstrate that colonic IECs are capable of producing estradiol while also responding to local estrogen via signaling through not only ERβ, but also ERα and GPER1.

### IEC-specific deletion of ERα protects females against intestinal injury

Given our discovery that ERα is expressed in IECs, coupled with the critical function of IECs in maintaining the barrier between the gut microbiota and host, we hypothesized that the protective impact of global ERα deletion (13) was primarily dependent upon changes in IEC function. Therefore, we generated a novel strain of ERα conditional knockout mice that express flanking LoxP sites under control of the villin promoter (ERα^fl/fl^/Villin^cre/+^, “ERα-cKO”, “cKO”). cKO mice appeared healthy with no apparent reproductive or developmental defects.

We challenged ERα-cKO males and females, as well as ERα-expressing (ERα^fl/fl^/Villin^+/+^, “Ctrl”) littermate controls, with 2.5% DSS for 5 days and evaluated intestinal injury on day 6 (Figure 2). As previously reported by us (13) and others (23), Ctrl female (Ctrl-F) mice exhibited less severe weight loss in response to DSS at day 6 than Ctrl-M (Figure 2A). Similar to our previous results in global ERα-KO mice (13), cKO-F mice lost less body weight compared to cKO-M mice and Ctrl-M and -F (green dotted line, Figure 2A). cKO-M mice lost more body weight than any other group (purple dotted line, Figure 2A). Interestingly, males and females responded differently to deletion of ERα, with cKO-M mice exhibiting more severe weight loss compared to WT-M (dotted versus solid purple lines, Figure 2A), but cKO-F mice maintaining weight compared to WT-F (dotted versus solid green lines, Figure 2A). Disease activity indices (DAI), comprising weight loss scores, stool consistency, and presence of fecal blood, revealed significant protection among cKO-F mice (Figure 2B).

**Figure 2:**
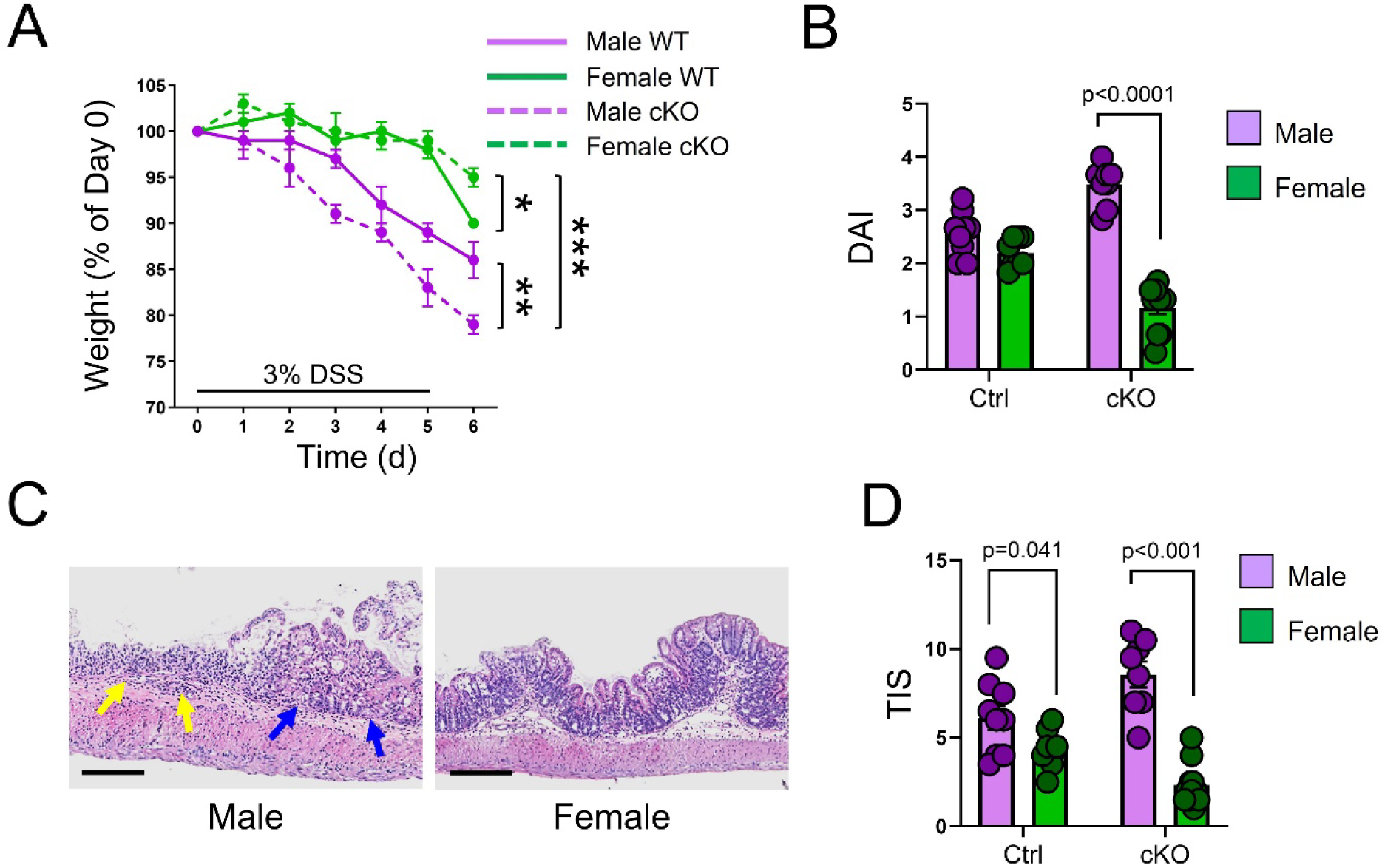
Intestinal epithelial-specific deletion of ERα protects females against intestinal injury. *(A)* Body weight loss in Ctrl (Villin^+/+^/ERα^flox/flox^) or ERα cKO (Villin^Cre/+^/ERα^flox/flox^) mice treated with 3% DSS for 5 days, changed to standard H2O for 1 day, and euthanized on day 6. Error bars represent SEM. *(B)* Disease activity index (DAI) scores for DSS-induced intestinal injury in Ctrl and cKO mice. *(C)* Representative hematoxylin and eosin-stained colon tissues collected from DSS-treated mice on day 6. Images were acquired with 20X magnification and scale bars = 10µm. *(D)* Total inflammation scores (TIS) based on histological inflammation of colon tissues collected from DSS-treated mice on day 6. For all figures, statistical analysis was performed with 2-way analysis of variance (ANOVA) and Tukey post hoc test. Individual points represent individual animals.

H&E-stained colon tissues from male and female cKO mice demonstrated significant epithelial erosion in cKO-M, along with increased inflammatory infiltrates (yellow arrows, Figure 2C) and changes in crypt architecture (blue arrows, Figure 2C). Consistent with this, total inflammatory scores (TIS) based on histological findings revealed enhanced inflammation in cKO-M colons compared to cKO-F and Ctrl colons (Figure 2D).

These findings directly support our hypothesis that IEC-specific ERα-mediated signaling contributes to increased barrier dysfunction and inflammation in a sex-specific fashion, and that removal of ERα signaling promotes protection from DSS-induced intestinal injury in females.

### Deletion of ERα or ERβ result in sex-specific changes to IEC lineage commitment

In order to determine the impact of ERα and ERβ-specific signaling on colonic IEC differentiation and function, we turned to the global knockout mice. The colonic epithelium of adult (4 to 12 months old) global ERβ-KO mice exhibits increased proliferation, decreased apoptosis, and reduced expression of differentiation-associated genes *Plec* (encoding plectin), *Ctnna1* (alpha-catenin), and *Krt20* (cytokeratin 20) (11). However, this earlier study did not report the sexes of experimental animals. In light of our discovery that colonic IECs also express ERα (Figure 1) and that its expression differentially impacts intestinal pathology upon DSS challenge (Figure 2), we hypothesized that ERα signaling in the absence of ERβ in ERβ-KO mice was the driver of these cellular changes. Thus, we compared colon morphology and inflammation in young (8 to 12 weeks old) WT, ERα-KO, and ERβ-KO mice.

H&E-stained colon tissues showed no evidence of altered colonic architecture in any of the genotypes, although female ERα-KO and ERβ-KO colons displayed moderate mucosal thickening compared to WT controls (yellow bars, Figure 3A) and male ERα-KO colons exhibited moderate goblet cell hyperplasia (yellow arrows, Figure 3A). As defined by the presence of polymorphonuclear cells or mononuclear cells, active and chronic inflammation, respectively, was not observed in any of the colon tissues (Figure 3B).

**Figure 3:**
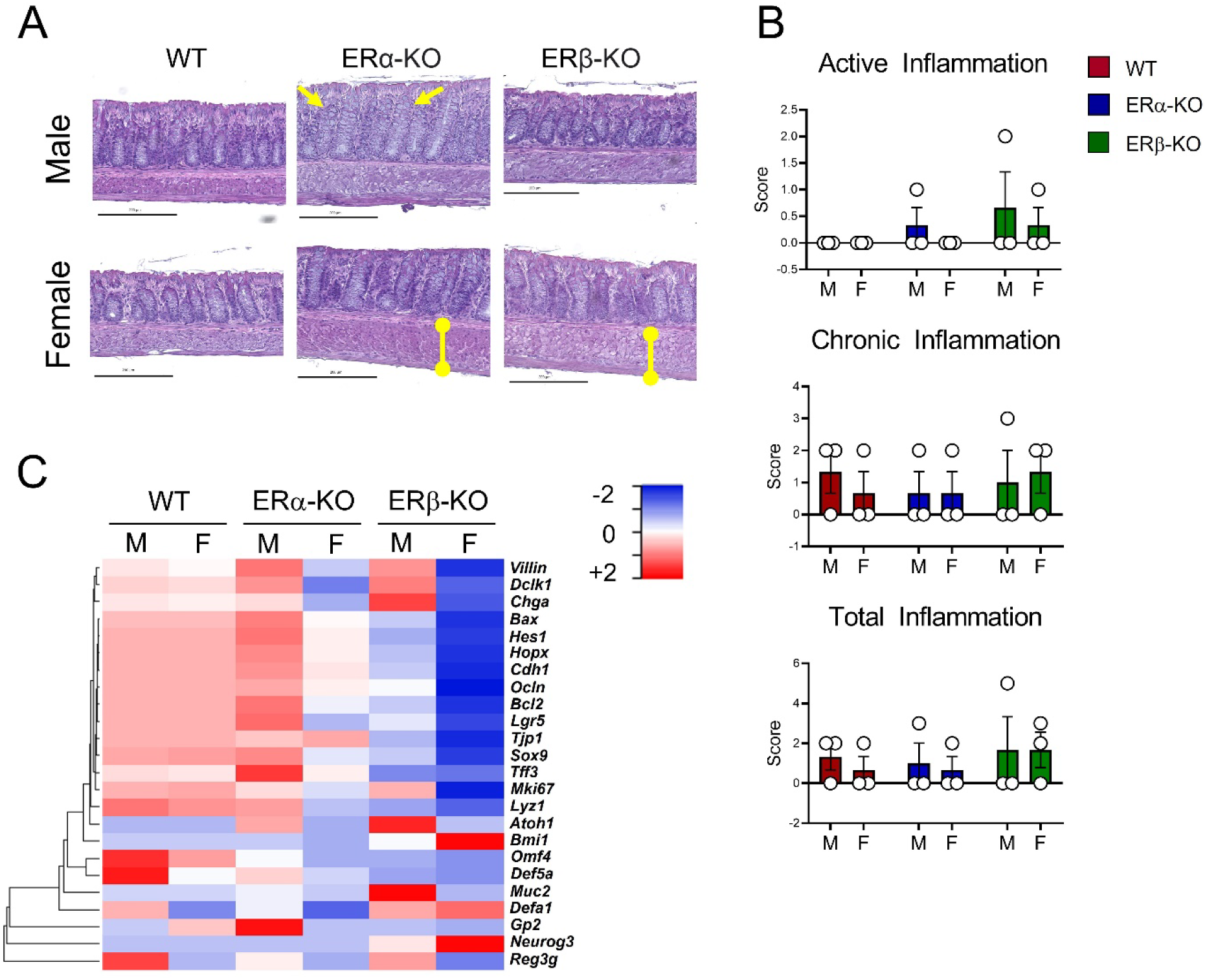
Deletion of ERα or ERβ alters expression of intestinal epithelial lineage genes. *(A)* Representative images of H&E-stained colon tissues from 8-10 week old WT, ERα-KO, and ERβ-KO mice. Scale bar = 200um. *(B)* Histopathological scoring of active, chronic, and total inflammation observed in H&E-stained colon tissues. Each dot represents an individual mouse. *(B)* mRNA was isolated from full-thickness colon tissues of 8-10 week old WT, ERα-KO and ERβ-KO mice. qPCR was performed for indicated genes relevant for intestinal epithelial lineage commitment.

Next, full-thickness colon tissues were screened for the expression of genes implicated in IEC lineage differentiation to determine whether ERα or ERβ contributes to colonocyte lineage specification. qPCR of *Lgr5*, *Bmi1*, *Hopx*, *Neurog3*, *Villin*, *Defa1*, *Chga*, and other markers showed similar expression between WT-M and -F tissues (Figure 3C, left two columns) for most genes, with only three showing differences between male and female samples. Specifically, *Defa1* and *Reg3g* were higher in male compared to female whereas *Gp2* was higher in female compared to male. Three genes (*Villin*, *Dclk1*, and *Chga*) showed similar sex-specific expression patterns in ERα-KO and ERβ-KO samples, with upregulation in male samples compared to female, suggesting that there is redundancy in ERα- and ERβ-specific regulation of these genes. In general, ERα-KO samples (Figure 3C, middle two columns) showed the most significant sex-specific changes, with male samples showing upregulation of many lineage genes and females showing downregulation. In contrast, both male and female ERβ-KO samples (Figure 3C, right two columns) were significantly downregulated for nearly every lineage gene, suggesting that ERα signaling in the absence of ERβ fails to maintain proper lineage specification of colonocytes, including *Lgr5*- and *Bmi1*-expressing stem cell populations.

We also asked whether the remaining estrogen receptors show compensatory expression in response to deletion of ERα or ERβ. Colonic IECs were assessed for mRNA expression of *Esr1* (for WT and ERβ-KO samples), *Esr2* (for WT and ERα-KO) samples, and *Gper1* (for WT, ERα-KO, and ERβ-KO samples). We found no significant changes in gene expression of *Esr1* or *Esr2* (Supplemental Figure S2A-B). *Gper1* expression was reduced by ∼60% in male ERα-KO IECs compared to female, and consistent in all other cohorts (Supplemental Figure S2C). In addition, ERα-KO and ERβ-KO colonocytes express comparable levels of Cyp19A1 as do WT cells (Supplemental Figure S3A-B).

These findings are consistent with our observations that ERα-specific signaling in IECs contributes to a weakened barrier and increased susceptibility to inflammatory stimuli and provide a potential mechanism for the protective effects of ERα deletion in females (Figure 2).

### Colonoids derived from male ERα-KO cells display accelerated growth kinetics

Given the clear importance of the *Esr1* to *Esr2* expression ratio, we next sought to determine the impact of intestinal injury on ERα and ERβ expression. We challenged 8 to 12 week old WT mice (male and female) for 5 days with 3% DSS, followed by one day of regular drinking water, and then evaluated expression of *Esr1* and *Esr2* in colonic IECs. DSS treatment resulted in upregulation of *Esr1* in male IECs, leading to a significantly higher ratio of *Esr1*:*Esr2* in males compared to females (Figure 4A). These findings are consistent with the more severe impact of DSS observed in males (Figure 2) and a pathologic role for ERα (*Esr1*) signaling in barrier function and disease overall. Furthermore, our data suggest that the ratio of *Esr1* to *Esr2* is critical not only for colonocyte lineage specification (Figure 3), but potentially also IEC proliferation and differentiation.

**Figure 4:**
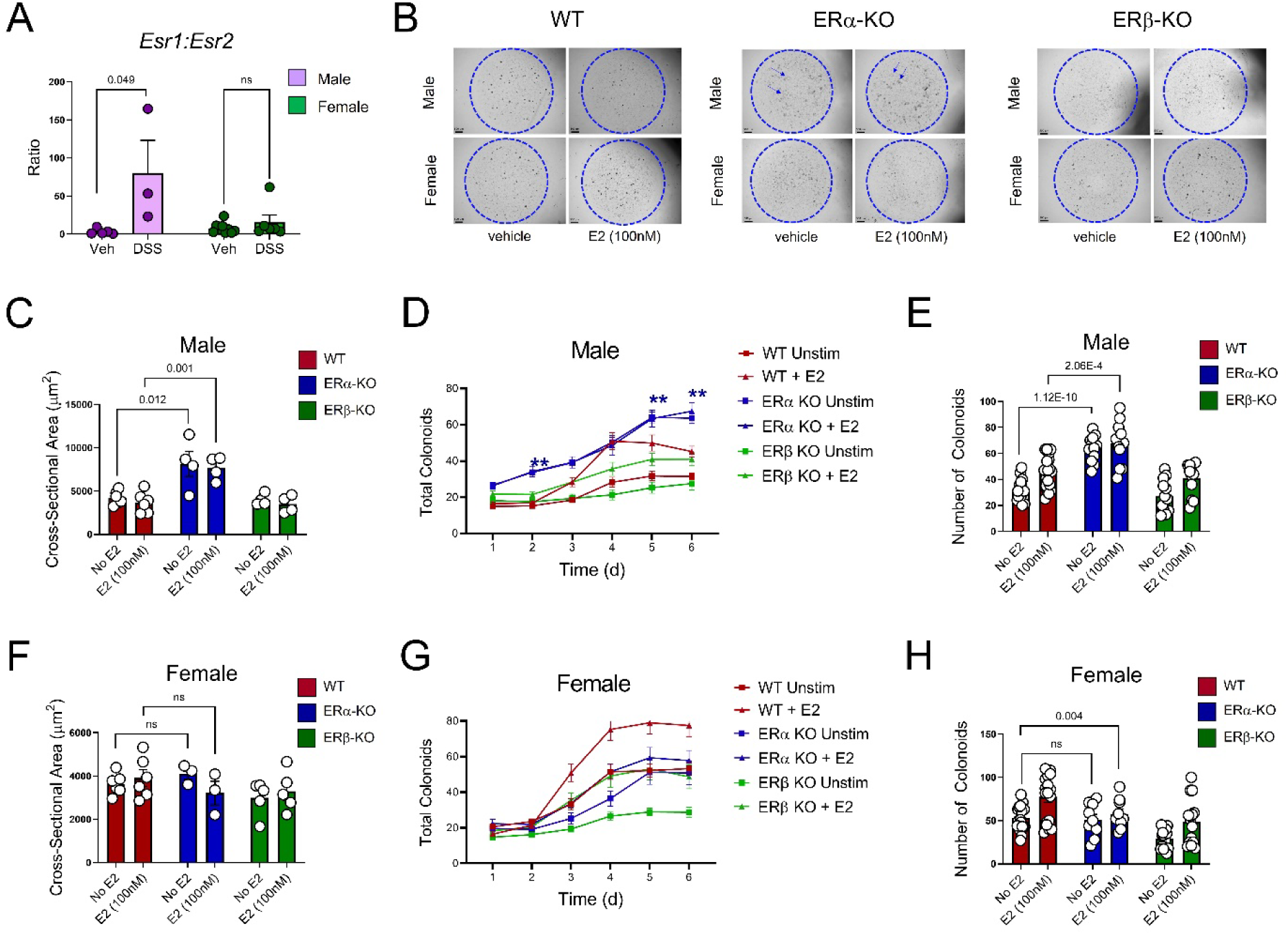
Colonoids derived from male ERα-KO cells display accelerated growth kinetics. *(A)* Primary colonocytes were isolated from male and female mice +/- DSS treatment and assayed for mRNA expression of *Esr1:Esr2* by qPCR. *(B)* Representative images of 3D colon organoids (colonoids) after 6 days in culture. Arrows point to enlarged spherical structures in male ERα-KO colonoids. *(C)* Cross-sectional area of day 6 male colonoids was calculated using ImageJ. *(D)* Number of distinct male colonoid structures was quantified on days 1-6. *(E)* Total number of male colonoids on day 6 is plotted for individual cultures. *(F-H)* Corresponding analyses for female colonoids.

To test this hypothesis, we established three-dimensional organoid models from colonic crypt cells of ERα-KO, ERβ-KO, and WT mice. 50 crypts were seeded per Matrigel dome and allowed to grow in the presence or absence of estrogen. Importantly, the frequency of Lgr5-expressing stem cells within each crypt preparation was comparable, with an average of 20-30% of EpCAM-expressing cells co-expressing Lgr5 (Supplemental Figure S4).

Imaging of colonoids on day 6 showed comparable morphology in WT and ERβ-KO cultures, with notably larger spherical structures in male ERα-KO cultures (blue arrows, Figure 4B). Images were taken of each Matrigel dome for days 1-6 of culture, and ImageJ software was used to quantify the size and quantity of structures in male (Figure 4C-E) and female (Figure 4F-H) cultures. Colonoids derived from male ERα-KO crypt cells were consistently larger than all other cohorts, with increased cross-sectional area (Figure 4C) and quantity (Figure 4D-E) compared to WT and ERβ-KO colonoids. Interestingly, the pro-proliferative effect of ERα-KO in male colonoids was observed with and without addition of exogenous estrogen, suggesting that ERα functions to suppress proliferation in an estrogen-independent manner.

### ERα-KO colonoids display sex-specific patterns of gene expression

To determine the mechanism(s) by which deletion of ERα impacts colonocyte function differently in males and females, we assayed day 6 colonoid samples for expression of several genes critical for IEC function, including tight junction genes, *Plec*, *Krt20*, and others (Figure 5). We observed striking, sex-dependent differences in gene expression among ERα-KO versus ERβ-KO colonoids, both in cultures not supplemented with estrogen (Figure 5A) and those supplemented with estrogen (Figure 5B). In particular, male ERα-KO colonoid samples showed significant downregulation of several functional genes, including *Ptk6*, *Cldn12*, *Krt20*, *Plec*, *Tjp1*, *Prkca*, *Ocln*, *Cldn3*, and *Cldn8*, in estrogen-supplemented cultures as well as non-supplemented cultures. In contrast, female ERα-KO cultures showed enhanced expression of most genes, including *Cldn3*, *Cldn8*, *Ocln*, *Tjp1*, and *Plec*. We re-analyzed the data to calculate ratios of gene expression for supplemented versus non-supplemented cultures and found that estrogen signaling altered the expression of IEC functional genes in sex-specific patterns (Figure 5C). The most significant changes in gene expression based on Z-scores were observed for male and female ERα-KO colonoids. Colonoid expression of remaining estrogen receptors (*Esr1* for ERβ-KO cultures, *Esr2* for ERα-KO cultures, and Gper1 for all cultures) was not significantly different for any group (Supplemental Figure S5), indicating that compensatory changes to receptor expression is not responsible for the changes observed in knockout colonoids.

**Figure 5:**
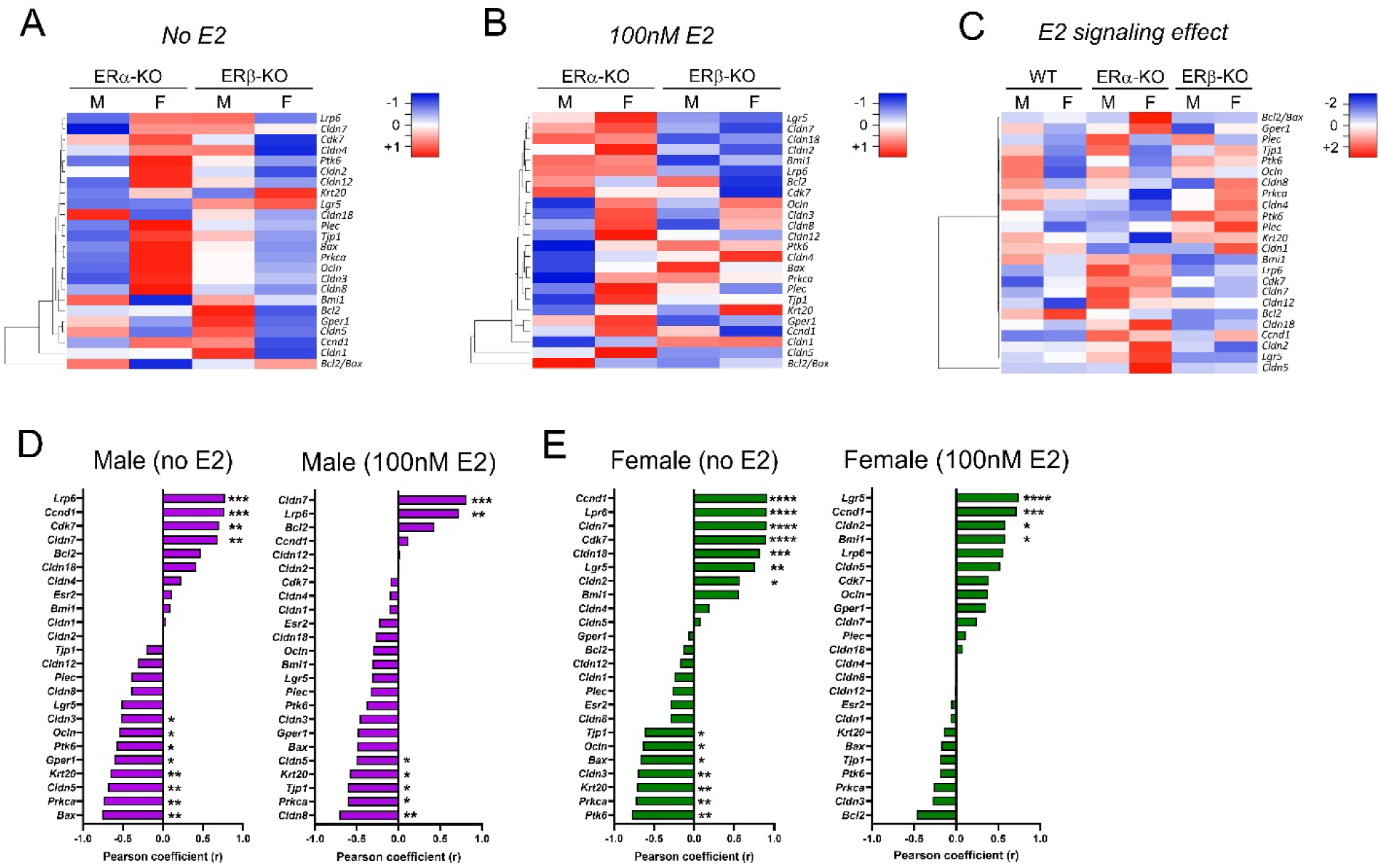
ERα-KO colonoids display sex-specific patterns of gene expression. *(A-C)* mRNA was prepared from day 6 colonoids and assayed by qPCR for expression of epithelial functional genes. Target gene expression was normalized to that of *Gapdh* and to expression within sex-matched, WT samples. *(A)* Relative expression of functional genes in colonoids not supplemented with E2. *(B)* Relative expression of functional genes in colonoids supplemented with 100nM E2. *(C)* Gene expression is shown for each column as a ratio of E2-supplemented colonoids/untreated colonoids. *(D)* Pearson correlation coefficients (r) were calculated for Esr1 versus each indicated gene in WT colonoids. * *p*<0.05; ** *p*<0.01; *** *p*<0.001; **** *p*<0.0001 (n=13-15/group).

Lastly, we asked how the expression of ERα correlates with that of IEC functional genes in WT colonoids. We calculated Pearson correlation coefficients for *Esr1* and each functional gene; data is plotted for male (Figure 5D) and female (Figure 5E) colonoids with and without estrogen. Genes showing significant correlations with *Esr1* included *Lrp6*, *Ccnd1*, *Cdk7*, and *Cldn7* (for both male and female), as well as *Cldn18*, *Lgr5*, and *Cldn2* (for female).

Collectively, our findings demonstrate that ERα-specific signaling results in sex-specific changes to IEC gene expression that underlie protection from challenge in females while enhancing susceptibility to intestinal injury in males.

## DISCUSSION

The intestinal epithelium represents one of the largest physical barriers separating environment from host tissue. Throughout the GI tract, a single cell layer of enterocytes functions as the gatekeeper between luminal contents, such as commensal microbes and dietary antigens, and mucosal host tissues. Thus, proper differentiation and function of intestinal epithelial cells (IECs) is critical for intestinal barrier function and host defense. IBD is characterized by defects in epithelial barrier function (24) including impaired IEC differentiation, loss of tight junction integrity, altered innate immune functions, and increased rates of apoptosis. Even minor, subtle changes to IEC function can have broad implications for overall barrier function, and studies in animal models have shown that loss of barrier function is an instigating event in IBD (25); therefore, it is imperative to understand how environmental signals impact IEC homeostasis and contribute to incomplete barrier function in settings of inflammation.

Sex differences have long been observed in human and experimental IBD (4, 8, 13, 26), leading to studies focused on the potential role of 17β-estradiol (E2) signaling on intestinal inflammation. Multiple cell types in the GI tract express estrogen receptors, including mucosal immune cells (26, 27) and colonic IECs (13). ERβ has been the focus of most of these studies, since colonic IECs are one of the strongest expressers of ERβ outside the female reproductive tract. Our previous work investigated the functional role of ERβ *in vivo*, specifically its ability to protect against DSS-induced intestinal injury. Although global deletion of ERβ did not significantly impact DSS susceptibility, deletion of ERα was found to be protective in female mice (13). This raised questions about the potential role of ERα in regulating sex-specific susceptibility to intestinal inflammation.

This study focused on the role of ERα-specific signaling in IECs, which represents an important and understudied aspect to understanding the impact of E2 signaling in the gut. Our results show that WT colonocytes express significantly higher levels of *Esr1* (encoding ERα) compared to *Esr2* (ERβ), indicating robust steady-state transcription of ERα. Although total protein levels of ERα and ERβ are comparable among male and female WT IECs, nuclear levels of ERβ are higher than those of ERα; this suggests that ERβ has a more significant role in regulating E2-dependent gene transcription under non-inflamed conditions. The precise role of ERβ homodimers in regulating transcription, compared to ERα homodimers or ERα/ERβ heterodimers, warrants further study.

Despite the enrichment of nuclear ERβ compared to ERα, our results reveal critical roles for ERα-specific signaling in IEC differentiation and function. Colon tissues from adult, unchallenged ERα-KO mice showed mild goblet cell hyperplasia, especially in males, indicating that ERα may normally function to maintain goblet cell differentiation and metabolic function. Transcriptional profiles of primary colonocytes also showed significant differences, with higher levels of *Tff3* (encoding Trefoil Factor 3) and *Gp2* (Glycoprotein 2) in IECs isolated from male ERα-KO mice compared to WT and ERβ-KO controls. Both *Tff3* and *Gp2* are broadly associated with maintenance of the mucosal barrier and epithelial repair, suggesting that ERα may limit the expression of these genes under homeostatic conditions in males. Interestingly, similar patterns of gene expression were not observed in IECs isolated from female ERα-KO mice, suggesting that ERα’s regulatory role on IEC differentiation and function is sex-specific.

Our studies in colonic organoids lend further support to the notion that ERα differentially impacts male and female colonocyte development. Colonoids derived from male ERα-KO mice grew consistently larger (greater surface area) and faster (more structures) than colonoids derived from female ERα-KO mice and ERβ-KO and WT controls, suggesting that ERα may function to restrain proliferation and differentiation of colonocytes in males. Interestingly, the enhanced growth in ERα-KO male colonoids was observed with or without addition of exogenous E2. This suggests that tonic expression of ERα normally functions to limit colonocyte proliferation. However, it is possible that in the absence of ERα (ERα-KO colonoids), there is altered activation of ERβ even without the addition of exogenous E2; for example, other steroid receptors have been shown to activate in response to kinase signaling, promiscuous binding by other ligands, or interactions with other transcription factors (reviewed in (28)). Un-liganded ERβ has been reported to induce expression and suppression of numerous genes normally associated with E2-mediated ERα signaling (29); therefore, our observations of enhanced colonocyte growth in ERα-KO colonoids may be due to either the direct loss of ERα, or altered activation of ERβ.

Interestingly, female colonoids showed a trend towards accelerated growth and differentiation in response to exogenous E2. That female colonocytes may be more responsive to E2 signaling suggests potential epigenetic regulation in response to heightened E2 signaling *in vivo*. Studies from breast cancer literature have revealed numerous epigenetic changes in response to E2 signaling versus deprivation (30), suggesting that elevated levels of E2 in women may function to alter the genetic landscape.

Our analysis of colonocyte-associated functional genes also revealed striking sex differences between ERα-KO and ERβ-KO colonoids. Female ERα-KO colonoids upregulated expression of most key functional genes, including *Ccnd1*, *Lrp6*, *Cldn7*, *Cdk7*, *Cldn18*, and *Lgr5*. In contrast, male ERa-KO colonoids showed only moderate upregulation of a more limited set of genes, including *Lrp6*, *Ccnd1*, and *Cdk7*. This general trend was observed both in E2-supplemented colonoid cultures and non-supplemented cultures, reinforcing earlier findings that the expression of ERα is a more significant contributor to colonocyte differentiation than E2-dependent signaling through this receptor.

Collectively, our results identify ERα as a critical regulator of sex-specific differences in colonocyte growth and differentiation. The significant reductions in IEC-related functional genes in ERα-KO males indicates that ERα plays an important role in maintaining expression of these functional genes under healthy conditions. We therefore propose that under homeostatic conditions, ERα normally functions to restrain proliferation and differentiation of male colonocytes, contributing to intestinal homeostasis. Future studies to better understand the impact of ERα downstream signaling in male versus female IECs are clearly warranted, given the identification of ERα as a central hub mediating sex differences in intestinal health and disease.

## MATERIALS AND METHODS

### Mice

Strains used include wild-type (WT) C57BL/6 (stock #000664, Jackson Labs, Bar Harbor, ME), global ERα-KO (stock #004744, Jackson Labs), global ERβ-KO (stock #004745, Jackson Labs), and intestinal epithelial cell-specific ERα-KO (ERα-CKO), generated via backcross of ERα-floxed mice (ERα^flox/flox^, gift of J.A. Gustafsson (31)) with Villin-cre-expressing mice (stock #021504). All experimental mice were bred and housed at CWRU under Specific Pathogen Free (SPF) conditions, fed standard laboratory chow (Harlan Teklad, Indianapolis, IN) and maintained on a 12-hour light/dark cycle. All animal procedures were approved by the Case Western Reserve University Institutional Care and Use Committee (protocol 2021-0014).

### DSS-Induced Intestinal Injury Model

8- to 12-week-old mice were supplemented with 3% colitis-grade DSS (36,000–50,000 daltons; MP Biomedicals, Solon, OH) dissolved in water and sterile-filtered through a 0.22um filter. Mice were evaluated for 6 days for weight loss, diarrhea, and rectal bleeding.

### Histologic Assessment of Colonic Inflammation

Mouse colon tissues were formalin-fixed, paraffin embedded, cut to 4µm, and stained with H&E as previously described (13). Intestinal damage was evaluated by a pathologist blinded to mouse genotype and sex using an established scoring system (13). Digital images were obtained using an Olympus VS120 slide scanner equipped with a 10× objective and 2/3-inch high sensitivity, high resolution CCD camera (Olympus Life Science).

### Intestinal Epithelial Cell Isolation

Colon tissues were harvested and flushed with ice-cold HBSS. Tissue was opened longitudinally, cut into small pieces, and transferred to 50mL conical tubes containing 15mL epithelial cell solution (HBSS containing 10nM Hepes, 10nM EDTA, 100U/mL Pen/Strep, 2% FBS, and 100ug/mL DNase I (32)). Tissues were incubated in a 37°C water bath for 15 minutes with gentle agitation every 5 minutes, then transferred to ice for 10 minutes. Tissues were then transferred to new tubes with fresh HBSS and shaken vigorously for 40 seconds. Resulting cells (containing crypt aggregates) were centrifuged and then resuspended using gentle pipetting in 5mL pre-warmed TrypLE containing 100ug/mL DNase I. Samples were incubated for 5 minutes, then gently pipetted for several more minutes until single cells were visible under a microscope. 1mL FBS was added to stop the reaction, then cells were strained through a 40um strainer and washed.

### Organoid Culture and Quantitation

Colonic crypt aggregates were prepared as described above. Crypts were suspended in Matrigel Matrix domes (Corning Life Sciences, Glendale, AZ) at a density of 50 crypts per 20uL Matrigel dome. Inverted domes were polymerized at 37°C for 30min, and then covered with Intesticult Organoid Growth Media/Mouse (StemCell Technologies, Vancouver, CA) and cultured for 1-6 days. Where indicated, estrogen receptor agonists were added to the Intesticult media every 24 hours at concentration of 100nM: 17β-estradiol (E2, endogeneous ER agonist, Tocris Bioscience); 4,4’,4’’-(4-Propyl-[1H]-pyrazole-1,3,5-triyl)trisphenol (PPT, ERα-selective agonist, Tocris Bioscience); or Diarylpropionitrile (DPN, ERβ-selective agonist, Tocris Bioscience). Organoids were imaged using a Keyence BZ-X810 inverted phase contrast microscope (Keyence, Chicago, IL) with a 10X PlanFluor_DL objective. Images were quantified using ImageJ software.

### RNA Isolation and Gene Expression Analysis

RNA was extracted from colon tissue samples using TRIzol (Thermo Fisher Scientific, Waltham, MA) according to the manufacturer’s instructions. RNA was extracted from primary intestinal epithelial cells and organoid cultures using the High Pure RNA Isolation Kit (Roche Life Science, Indianapolis, IN) according to the manufacturer’s instructions. Total RNA was quantified using a NanoDrop Lite spectrophotometer (Thermo Fisher) and 1µg of RNA was reverse-transcribed to cDNA using the High-Capacity cDNA Reverse Transcription Kit (Thermo Fisher). Real-time qPCR was performed using Taqman gene-expression assays (Thermo Fisher) on an Applied Biosystems QuantStudio 3 real-time PCR system. Gene-expression values were normalized to those of β2-microglobulin or GAPDH (housekeeping genes) and fold-change values were calculated using the ΔΔCT method.

### Protein Isolation, Western blot, and Densitometry

Colon tissues were homogenized using a bead beater. Tissue homogenates or epithelial cells were lysed in RIPA buffer (for total protein) or NE-PER extraction kit (Thermo Fisher, for nuclear/cytoplasmic protein) according to the manufacturer’s instructions. All lysis buffers contained protease/phosphatase inhibitor (Thermo Fisher). Protein concentrations were determined using Pierce BCA Protein Assay (Thermo Fisher) and equivalent amounts were loaded on NuPage Bis/Tris gels (Thermo Fisher) and blotted for indicated proteins. Antibodies used for western blots included α-Cyp19A1 (NSJ Bioreagents #RQ4643) α-ERα (Novus #NB300-560), α-ERβ (Novus #NB120-3577), α-GPER1 (Abcam #ab260033), α-Lamin B1 (Cell Signaling #12586), and α-β-Actin (Cell Signaling #12262).

### Data Analysis and Statistics

Graphical analysis was performed using GraphPad Prism 10 (GraphPad Software, La Jolla, CA). Statistics were performed using ANOVA one-way comparisons with Tukey’s post-hoc tests. P values ≤ 0.05 were considered significant.

## Supplemental Figures and Figure Legends

**Supplemental Figure S1:**
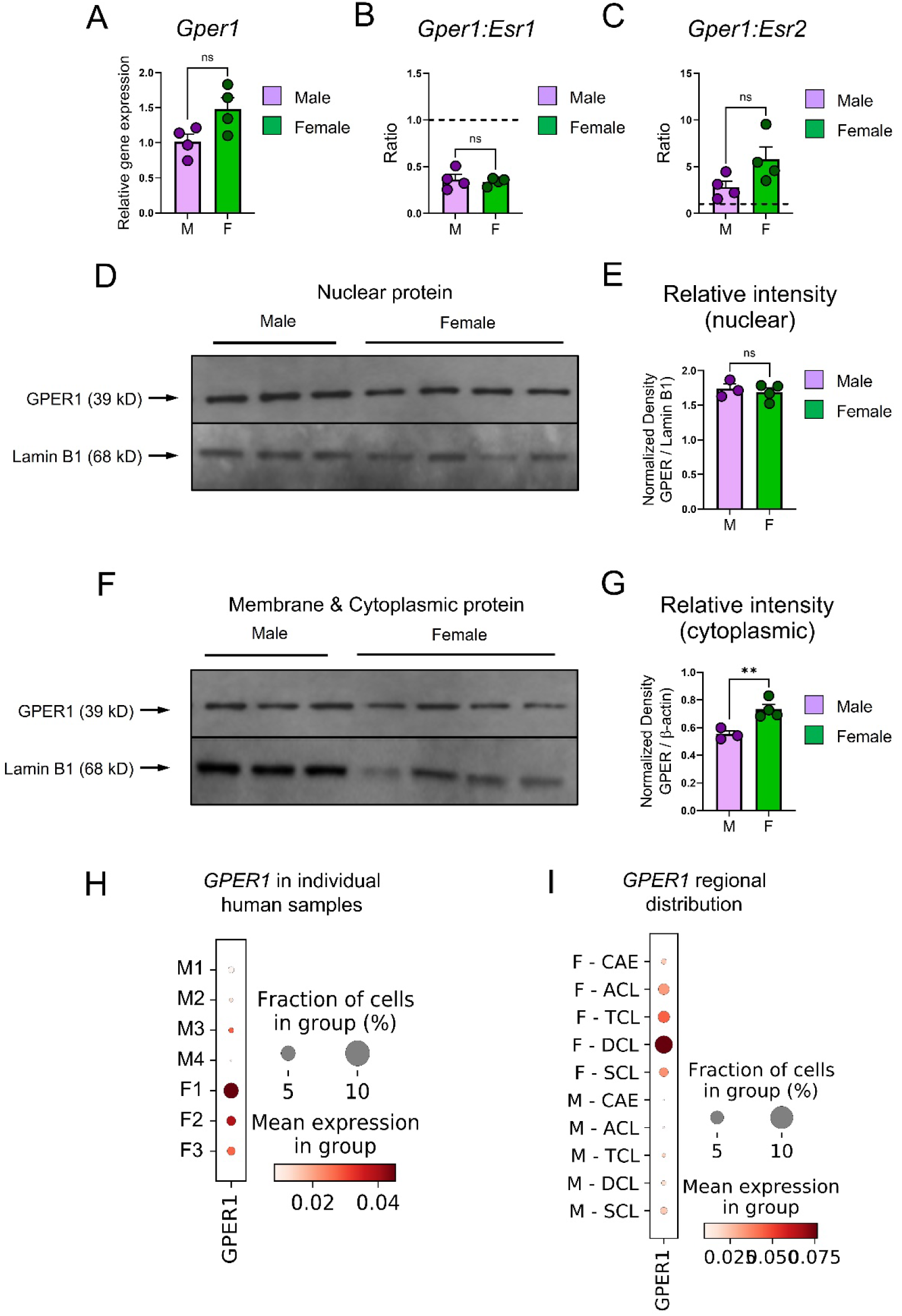
The g-protein coupled estrogen receptor GPER1 is expressed by murine and human intestinal epithelial cells. *(A)* mRNA expression of *Gper1* in primary colonocytes was determined by qPCR and normalized to expression of *Gapdh*. *(B-C)* Ratios of *(B) Gper1:Esr1* and *(C) Gper1:Esr2* gene expression were calculated for male and female colonocyte samples. *(D)* Protein expression of Gper1 was determined in nuclear lysates from primary colonocytes and *(E)* relative intensity was calculated using densitometry. *(F)* Protein expression of Gper1 was determined in cell membrane/cytoplasmic lysates from primary colonocytes and *(G)* relative intensity was calculated using densitometry. *(H-I)* mRNA expression of GPER1 in healthy adult human colonocytes was assessed using a publicly-available single-cell RNA-Seq dataset. *(H)* Expression of GPER1 in male (M) versus female (F) colonocytes. *(I)* Expression of GPER1 in colonocytes isolated from indicated regions of the colon. For all figures, statistical analysis was performed using a student’s t-test. Individual points represent individual animals. For all figures, statistical analysis was performed with 2-way analysis of variance (ANOVA) and Tukey post hoc test. Individual points represent individual animals.

**Supplemental Figure S2:**
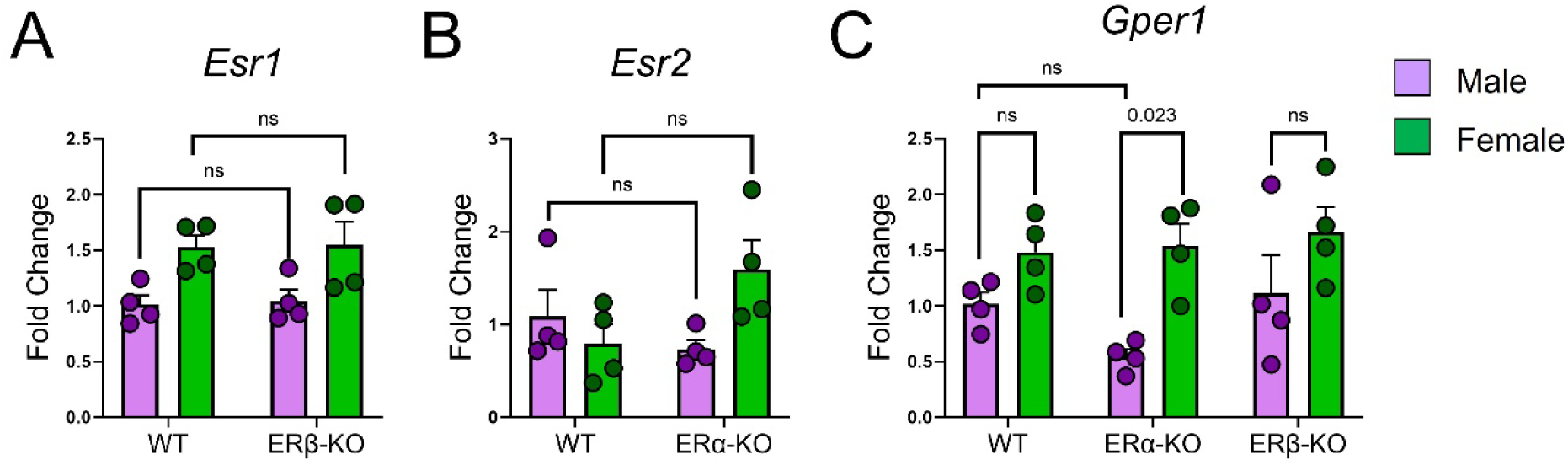
Expression of remaining estrogen receptors is consistent in intestinal epithelial cells isolated from ERα-KO and ERβ-KO mice. *(A-C)* Intestinal epithelial cells were isolated from colons of 8-10 week old WT, ERα-KO, and ERβ-KO mice. Gene expression of *(A) Esr1*, *(B) Esr2*, and *(C) Gper1* was determined by qPCR. For all figures, statistical analysis was performed with 2-way analysis of variance (ANOVA) and Tukey post hoc test. Individual points represent individual animals.

**Supplemental Figure S3:**
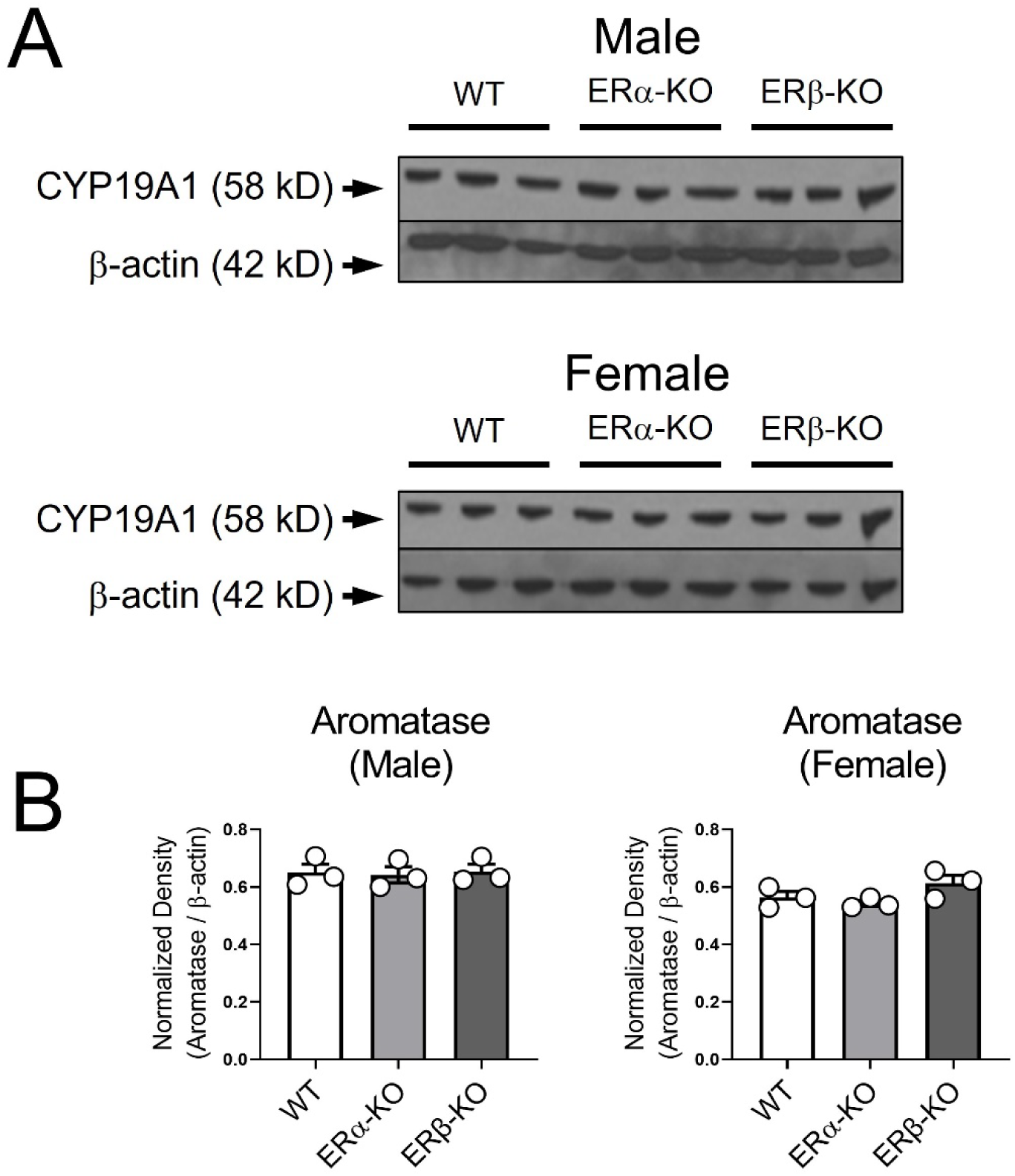
ERα-KO and ERβ-KO IECs express normal levels of aromatase. Primary intestinal epithelial cells were isolated from colons of 8-10 week old WT, ERα-KO, and ERβ-KO mice. Whole cell protein lysates were prepared and immunoblotted. *(A)* Representative blots showing expression of Cyp19A1 in primary colonocytes. *(B)* Relative intensity of Cyp19A1 expression was determined by densitometry.

**Supplemental Figure S4:**
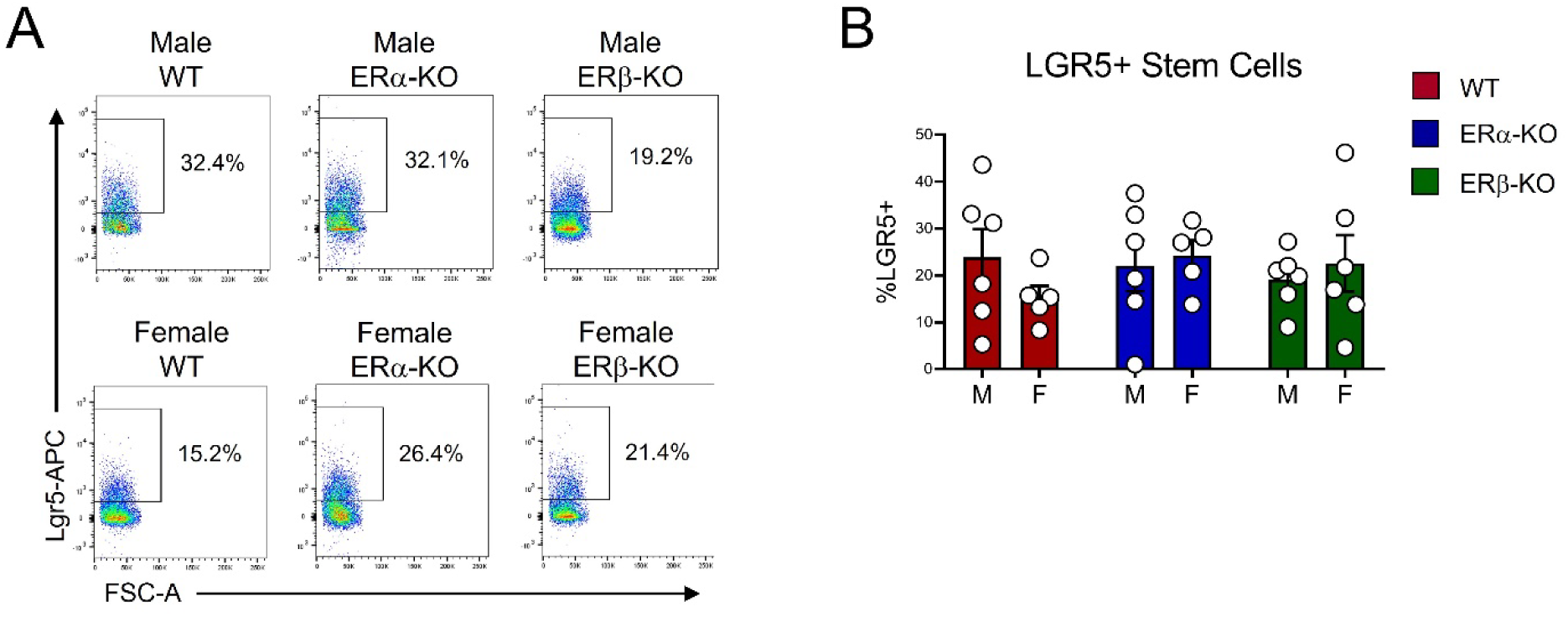
Intestinal crypts of WT, ERα-KO and ERβ-KO mice contain comparable expression of Lgr5+ stem cells. Primary intestinal epithelial cells were isolated from colons of 8-10 week old WT, ERα-KO, and ERβ-KO mice. Cells were analyzed for expression of Lgr5 by flow cytometry. *(A)* Representative histograms showing flow cytometric staining of Lgr5 within the live, Epcam+ population. *(B)* Quantification of flow cytometric staining. Each dot represents one mouse.

**Supplemental Figure S5:**
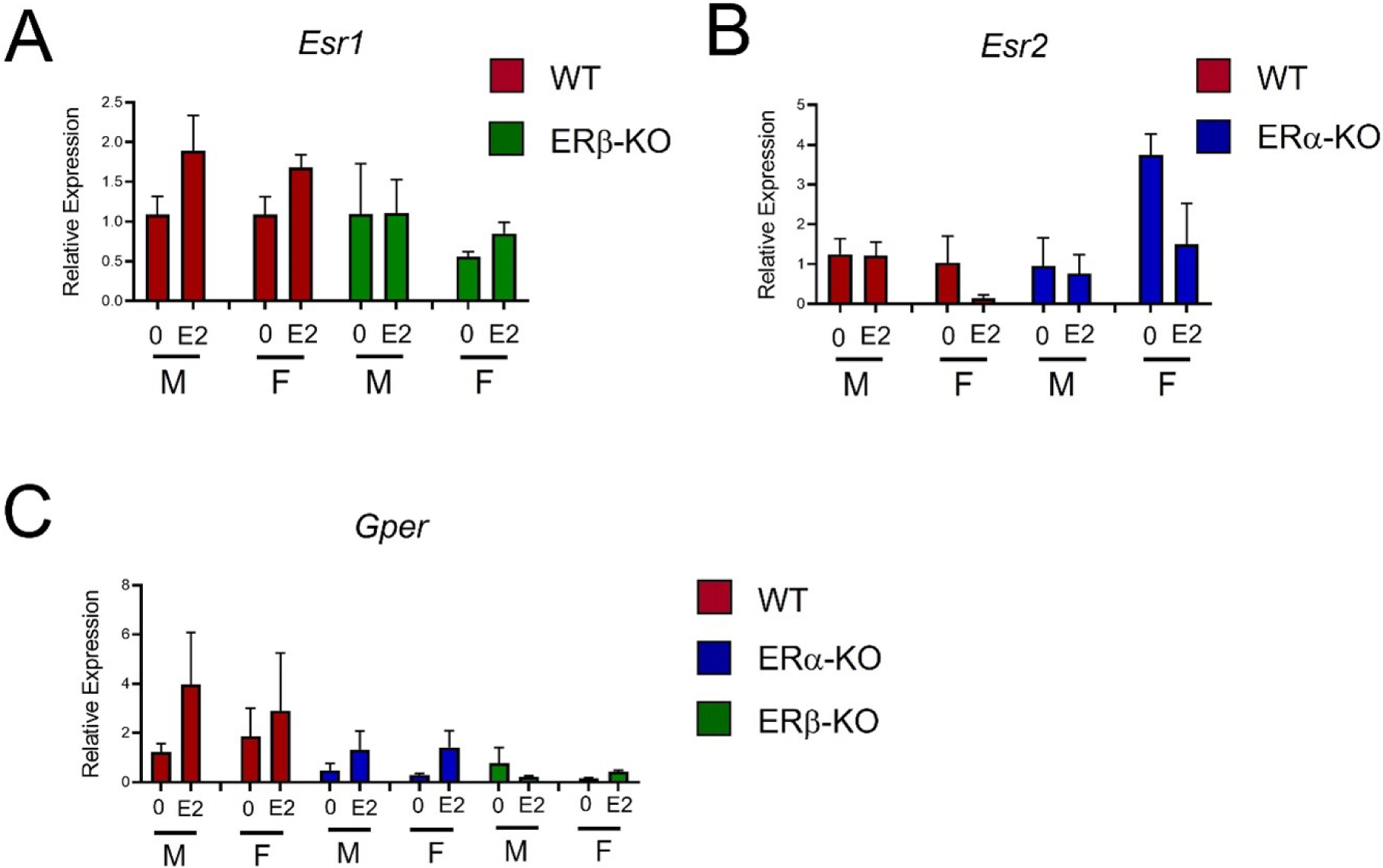
Colonoids do not display significant differences in gene expression of estrogen receptors. mRNA was isolated from day 6 colonoids and assayed for expression of *(A) Esr1*, *(B) Esr2*, and *(C) Gper1* by qPCR. Expression was normalized to that of *Gapdh*. Data represents fold change relative to WT male colonoids (no E2) and bars display mean +/- SEM.

## Grant Support

R01DK128143 (W.A.G.)

CWRU DDRCC support through funding from the National Institutes of Health, P30 DK097948 Crohn’s and Colitis Foundation Senior Research Award # 1169305 (W.A.G.)

R01DK115806 and R01DK109559 (S.T.M.)

UNC/NCSU CGIBD support through funding from the National Institutes of Health, P30 DK034987

F31DK136305 (K.A.B)

## Data Statement

All data are available upon request.

## Author Contributions

Guillermo Pereda: Writing – original draft, review and editing, Data curation, Investigation, Formal Analysis

Adrian Kocinski: Writing – review and editing, Data curation, Investigation, formal analysis Alyssia Broncano: Investigation

Sarah McNeer: Investigation Michelle Raymond: Investigation Nicholas Ziats: Investigation

Keith Breau: Investigation, Formal Analysis Joseph Burclaff: Investigation, Formal Analysis Scott Magness: Investigation

Wendy Goodman: Conceptualization, Funding Acquisition, Data curation, Writing – original draft, Writing – review and editing, Supervision, Project Administration

**<u>Article Synopsis</u>:** Estrogen receptor alpha (ERα) was identified as a key regulator of sex differences in response to intestinal injury. Studies using epithelial-specific ERα-knockout mice revealed critical protective functions for ERα in the proliferation, differentiation, and function of male colonocytes.

## Abbreviations

CD: Crohn’s disease
E2: estrogen
ERα: estrogen receptor alpha
ERβ: estrogen receptor beta
GPER1: g-protein coupled estrogen receptor 1
IBD: inflammatory bowel disease
IEC: intestinal epithelial cell
UC: ulcerative colitis

